# Automatic Abdominal Multi Organ Segmentation using Residual UNet

**DOI:** 10.1101/2023.02.15.528755

**Authors:** Gowtham Krishnan Murugesan, Diana McCrumb, Eric Brunner, Jithendra Kumar, Rahul Soni, Vasily Grigorash, Anthony Chang, Anderson Peck, Jeff VanOss, Stephen Moore

## Abstract

Automated segmentation of abdominal organs plays an important role in supporting computer-assisted diagnosis, radiotherapy, biomarker extraction, surgery navigation, and treatment planning. Segmenting multiple abdominal organs using a single algorithm would improve model development efficiency and accelerate model deployment into clinical workflows. To achieve broadly generalized performance, we trained a residual UNet using 500 CT/MRI scans collected from multi-center, multi-vendor, multi-phase, multi-disease patients, each with voxel-level annotation of 15 abdominal organs. Using the model trained on multimodality (CT/MRI), we achieved an average dice of 0.8990 in the held-out test dataset with only CT scans (N=100). An average dice of 0.8948 was achieved in the held-out test dataset with both CT and MRI scans (N=120. Our results demonstrate broad generalization of the model.

## 1 Introduction

Abdominal organ segmentation in Computed Tomography (CT) scans is an essential step for disease diagnosis, radiotherapy planning, surgery navigation and biomarkcr cxtraction[6][2][5]. Manual annotation is tedious, time consuming, and prone to both inter and intra-observer variability[8]. In addition, the presence of artifacts, heterogenous lesions, the variation between patients, and different scanners can lead to significant variances in organ appearance and makes it a complex problem for annotation[6]. This leads to leads to inter/intra-observer variabilities in msnusl annotations due to the low contrast of soft tissues in CT scans. Automating organ segmentation could complement clinical workflow in this aspect and accelerate computer aided diagnosis [2].

Inspired by the success of deep learning (DL) methods in image segmentation, especially for multi organ segmentation, researchers have proposed various methods for abdominal organ scgnicntation[1]. Several recent studies have reported state of art segmentation results on individually segmented abdominal organs such as the liver, kidneys, and the pancreas[5] [6] [9]. Gibson ct al presented a registration-free deep learning method in a cross-validation on a multi-center data set with 90 subjects to segment eight abdominal organs[2]. Turkay et al reported high qualitative and quantitative accuracy in segmenting liver, spleen, and kidney on healthy MRI data from UK Bio Bank [?]. However, most of the study contains dataset from single center and single disease cases on limited dataset, which make it unclear whether the models are broadly generalized in unseen diverse dataset [2].

To address the above drawbacks, Abdominal Multi Organ Segmentation (AMOS) challenge[4] provided a large-scale, clinical, and diverse abdominal multiorgan segmentation benchmark. AbdomentCT-1K dataset provides 1000 CT scans from 12 medical centers, multi-vendor, and multi disease to address the same drawbacks but only has an annotation for four organs, liver, spleen, kidney, and pancreas. The AMOS datasets provided multimodal (500 CT scans and 100 MRI) scans collected from multi-center, multi-vendor, multi-modality, multi-phase, multi-disease patients, each with voxel-level annotations of 15 abdominal organs (spleen, right kidney, left kidney, gallbladder, esophagus, liver, stomach, aorta, inferior vena cava, pancreas, right adrenal gland, left adrenal gland, duodenum, bladder, prostate/uterus). The main motivation for this challenge is to develop a single model that can absorb data to segment multiple organs from multi-center and multi disease facilitates the model to absorb multi center and multi disease data to accurately segment multiple abdominal organs and accelerates the deployment of the model in clinical setting[4]. In this paper, we utilized multimodal scans (CT/MRI) to segment multiple abdominal organs and participated in both Task-I and Task-II of the AMOS challenge. We trained a residual UNet model to segment all 15 organs at once using the CT/MRI scans and tested the same for both tasks instead of developing a model individually for each task.

## 2 Materials and Methods

### 2.1 Data and Preprocessing

#### Task 1

Segmentation of abdominal organs (CT only): Task 1 aims to comprehensively segment 15 abdominal organs across large-scale and great diversity CT scans, a total of 500 cases with annotations of 15 organs (spleen, right kidney, left kidney, gallbladder, esophagus, liver, stomach, aorta, inferior vena cava, pancreas, right adrenal gland, left adrenal gland, duodenum, bladder, prostate/uterus) (Fig. 1).

**Fig. 1.**
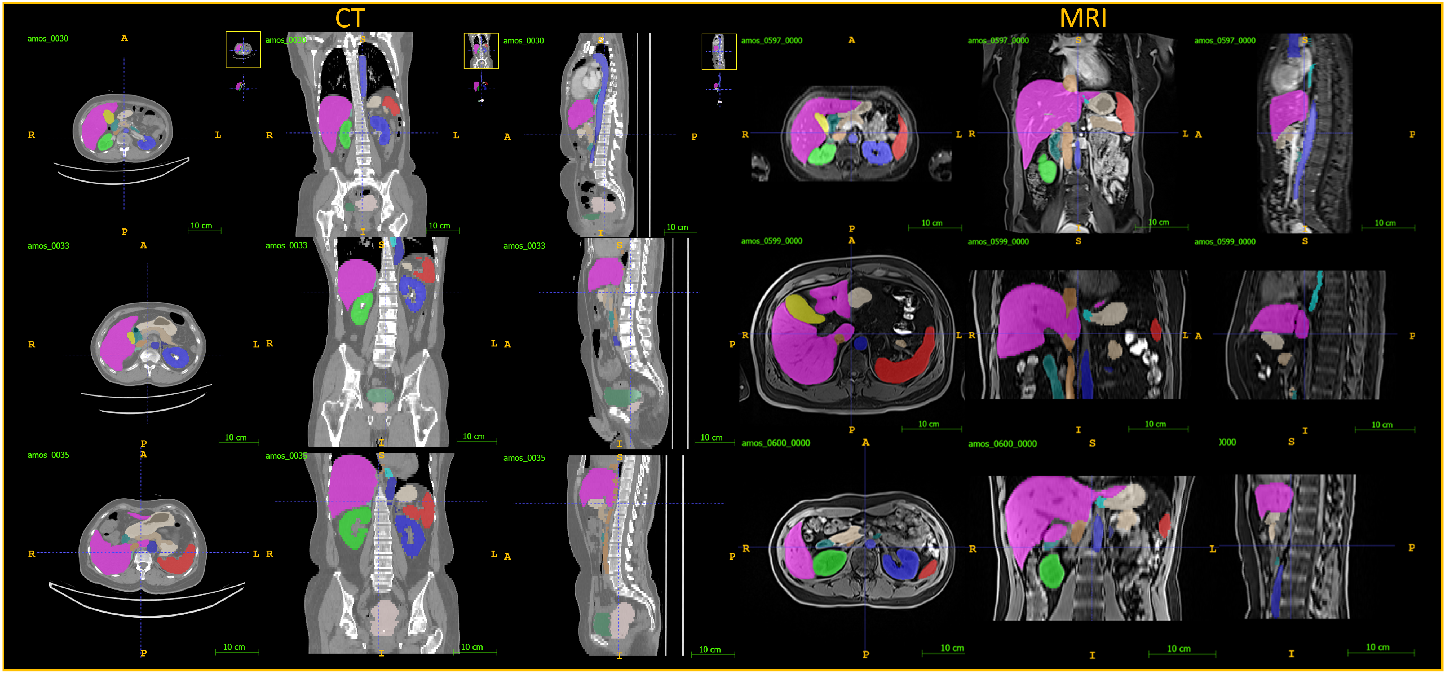
Representative CT and MRI scans provided by AMOS challenge with annotations for 15 abdominal organs

#### Task 2

Segmentation of abdominal organs (CT and MRI): this task extends the image modality target of Task 1 to the MRI modality. Task 2 aims to segment 15 abdominal organs, under such a “Cross Modality” setting, using a single algorithm to segment abdominal organs from both CT and MRI. Specifically, additional 100 MRI scans with the same type of annotation is provided.

We trained a model for Task 2 and utilized the same for Task 1 as it covers the scope of Task 1. For Task 2, The AMOS dataset provided 241 scans for training and 120 scans for held out testing each with manual annotations for 15 organs. The training dataset was further randomly split into training (80

### 2.2 Model Training Methodology

#### Model Architecture

The nnUNET pipeline has achieved top tier performance in multiple medical imaging segmentation competitions. Analysis of the nnUNET pipeline and model architecture has shown that different variations sometimes perform better than the baseline nnUNET architecture [7] [3]. From this, a standard variant model using residual connections was proposed for training (see Fig. 2 and 3). The input image size of 64×160×160 with one channel, CT is used as input. Input is resampled down five times by convolution blocks with strides of 2. On the decoder side, skip connections are used to concatenate the corresponding encoder layers to preserve spatial information. Instance normalization and leaky ReLU activation in the network layers were used. This architecture initially used 32 feature maps, which then doubled for each down sampling operation in the encoder (up to 1024 feature maps) and then halved for each transposed convolution in the decoder. The end of the decoder has the same spatial size as the input, followed by a 1×1×1 convolution into 1 channel and a SoftMax function. Models are trained for five folds with the loss function of Dice Sorensen Coefficient (DSC) in combination with weighted cross entropy loss were trained. To prevent overfitting augmentation techniques such as random rotations, random scaling, random elastic deformations, gamma, correction augmentation, mirroring, and elastic deformation, were adopted. Each of the five models was trained for 1000 epochs with a batch size of eight using SGD optimizer and a learning rate of 0.01. Dice Similarity Coefficient (DSC), and normalized surface dice (NSD), will be used to assess different aspects of the performance of the segmentation methods.

**Fig. 2.**
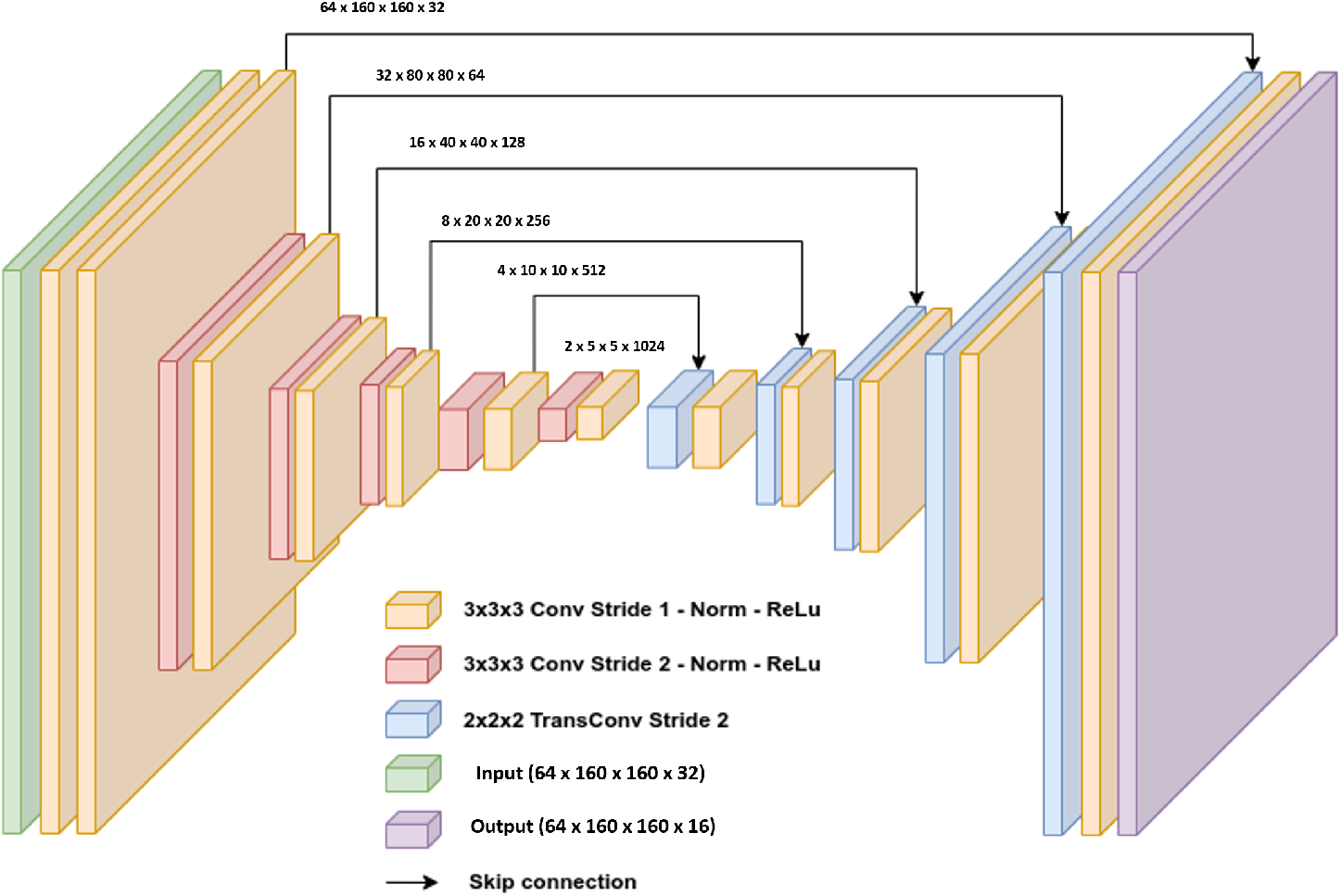
The layers of the UNET architecture used. The input is a volume of 64×160×160 with one channels, CT. Input is resampled down five times by convolution blocks with strides of 2. On the decoder side, skip connections are used to concatenate the corresponding encoder layers to preserve spatial information.

**Fig. 3.**
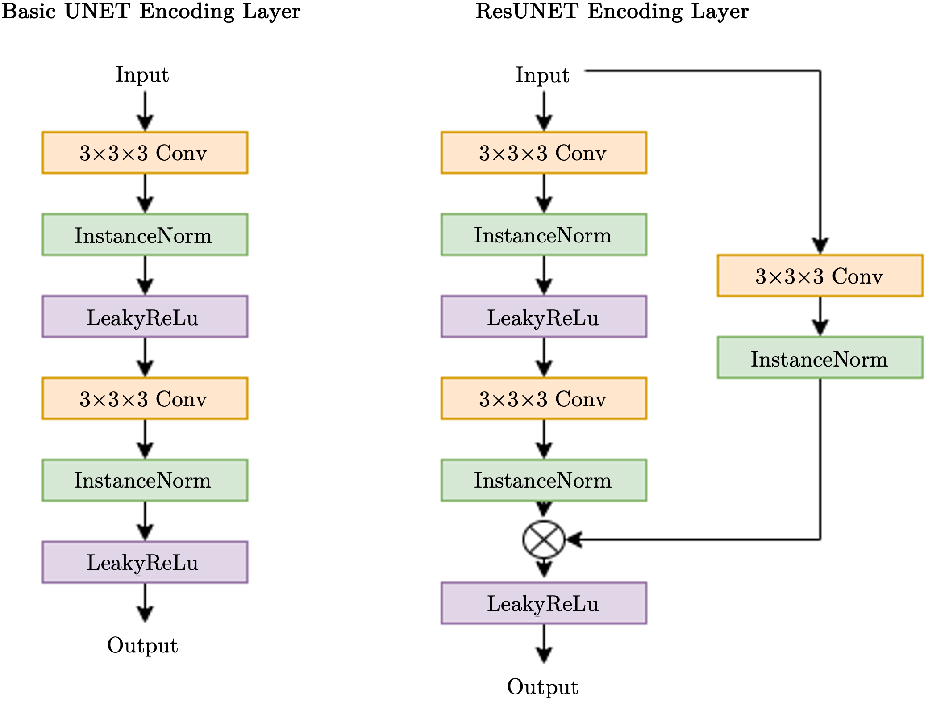
In one instance of our UNET models, each encoding layer is a series of Convolution, normalization, and activation function repeated twice. In another instance, ResUNET, each encoding layer adds a residual path with convolution and normalization.

### 2.3 Results

We trained a single residual UNet model for both tasks and our model achieved robust mean dice of 0.8990 and 0.8948 respectively for Task 1 with only CT scans and for Task II with both CT and MRI scans (Table.1)(Fig.4).

**Table 1.** Result on Task 1 (Only CT) and Task 2 (CT and MRI)

| Organ | Dice |  | NSD |  |
| --- | --- | --- | --- | --- |
|  | Task 1 (Only CT) | Task 2 (CT and MRI) | Task 1 (Only CT) | Task 2 (CT and MRI) |
| Spleen | 0.9701 | 0.9712 | 0.9125 | 0.9157 |
| Left Kidney | 0.9658 | 0.9666 | 0.9100 | 0.9120 |
| Right Kidney | 0.9656 | 0.9664 | 0.9060 | 0.9083 |
| Gallbladder | 0.8348 | 0.8295 | 0.7582 | 0.7537 |
| Esophagus | 0.8637 | 0.8574 | 0.7903 | 0.7930 |
| Liver | 0.9753 | 0.9761 | 0.8543 | 0.8591 |
| Stomach | 0.9302 | 0.9299 | 0.7957 | 0.7925 |
| Aorta | 0.9566 | 0.9553 | 0.9116 | 0.9147 |
| Postcava | 0.9219 | 0.9217 | 0.8093 | 0.8183 |
| Pancreas | 0.8840 | 0.8863 | 0.7505 | 0.7556 |
| Right Adrenal Gland | 0.8055 | 0.7869 | 0.8398 | 0.8312 |
| Left Adrenal Gland | 0.8163 | 0.8062 | 0.8346 | 0.8300 |
| Duodenum | 0.8427 | 0.8295 | 0.7167 | 0.7048 |
| Bladder | 0.8997 | 0.8907 | 0.7830 | 0.7752 |
| Prostate/Uterus | 0.8512 | 0.8427 | 0.6564 | 0.6498 |
| Mean | **0.8990** | **0.8948** | **0.8155** | **0.8169** |

**Fig. 4.**
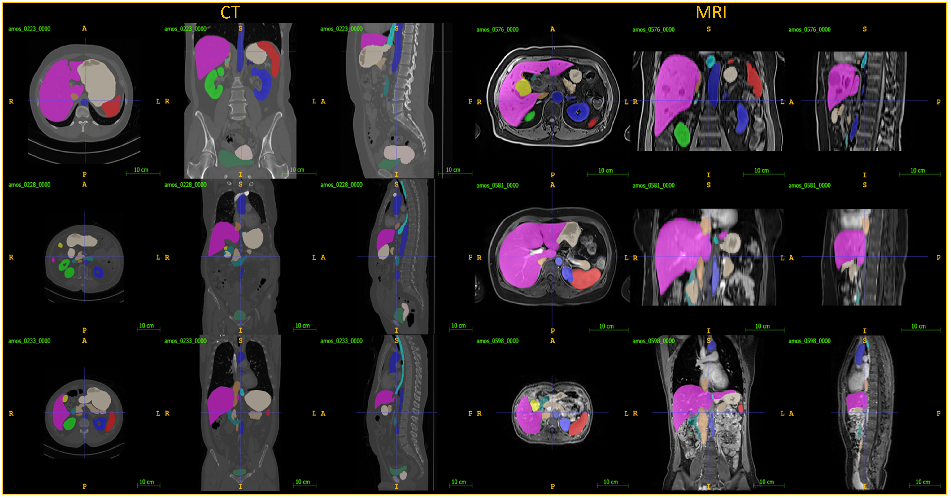
Prediciton on three CT and three MRI scans from held out test scans using our method

### 2.4 Discussion

Our method achieved similar performance in both tasks showing that it generalized well across both image domains (CT and MRI). Adding more folds, adaptive cnscmbling [7] and uncertainty aware segmentation correction may improve the segmentation performance.

### 2.5 Conclusion

We have trained a residual 3D UNet and achieved robust and generalized segmentation performance on cross domain image modalities. We achieved 7th rank in MICCAI AMOS 2022 challenge out of 48 teams participating in Task 2.

## Notes

### Competing Interest Statement

Employees of BAMF Health LLC

